# A realistic touch-transfer method reveals low risk of transmission for SARS-CoV-2 by contaminated coins and bank notes

**DOI:** 10.1101/2021.04.02.438182

**Authors:** Daniel Todt, Toni Luise Meister, Barbora Tamele, John Howes, Dajana Paulmann, Britta Becker, Florian H. Brill, Mark Wind, Jack Schijven, Baxolele Mhlekude, Christine Goffinet, Adalbert Krawczyk, Jörg Steinmann, Stephanie Pfaender, Yannick Brüggemann, Eike Steinmann

**Author notes:** **Corresponding author:** Prof. Dr. Eike Steinmann, Department of Molecular and Medical Virology, Ruhr University Bochum, Universitätsstr. 150, 44801 Bochum, Germany.

## Abstract

The current severe acute respiratory syndrome coronavirus 2 (SARS-CoV-2) pandemic has created a significant threat to global health. While respiratory aerosols or droplets are considered as the main route of human-to-human transmission, secretions expelled by infected individuals can also contaminate surfaces and objects, potentially creating the risk of fomite-based transmission. Consequently, frequently touched objects such as paper currency and coins have been suspected as a potential transmission vehicle. To assess the risk of SARS-CoV-2 transmission by banknotes and coins, we examined the stability of SARS-CoV-2 and bovine coronavirus (BCoV), as surrogate with lower biosafety restrictions, on these different means of payment and developed a touch transfer method to examine transfer efficiency from contaminated surfaces to skin. Although we observed prolonged virus stability, our results, including a novel touch transfer method, indicate that the transmission of SARS-CoV-2 via contaminated coins and banknotes is unlikely and requires high viral loads and a timely order of specific events.

## 1. Introduction

The current severe acute respiratory syndrome coronavirus 2 (SARS-CoV-2) pandemic has created a significant threat to global health. Since effective treatments and access to vaccines is still limited for the broad population in most countries, diligent attention on transmission-based precautions is essential to limit viral spread. In particular considering the emergence of novel SARS-CoV-2 variants with possibly greater risk of transmission [1,2]. According to current evidence, SARS-CoV-2 is mainly transmitted through respiratory droplets and aerosols exhaled from infected individuals [3]. Respiratory secretions or droplets expelled by infected individuals can potentially contaminate surfaces and objects (fomites) and have been shown to persist on inanimate surfaces for days under controlled laboratory conditions [4,5]. Therefore, a clinically significant risk of SARS-CoV-2 transmission by fomites has been assumed [6–8]. The COVID-19 pandemic intensified the decline in the transactional use of cash, partly due to reduced consumer spending, but also due to concerns about the risk of banknotes transmitting the virus. This was observed for either sides, the retailers’ as well as the customers [9]. Indeed, frequently touched objects such as banknotes and coins have been suspected to serve as transmission vehicle of various pathogenic bacteria, parasites, fungi and viruses including SARS-CoV-2 [10,11]. However, the conditions presented in various experimental studies frequently do not resemble real-life scenarios (e.g. large virus inoculums, small surface area) and thereby potentially exaggerating the risk of transmission of SARS-CoV-2 by fomites [12,13]. Although different viruses are readily exchanged between skin and surfaces, the fraction of virus transferred is dependent on multiple factors including virus species and surface material [14]. The efficiency of pathogen transfer from the fomite to hands is an important parameter to model its potential for transmission and to implement effective hygiene measures, while avoiding unnecessary measures [15]. However, the transfer of SARS-CoV-2 from surfaces to skin has not been analyzed systematically. Here, we examined the stability of SARS-CoV-2 and bovine coronavirus (BCoV) as surrogate on different means of payment. We further implemented a new protocol to study the touch transfer efficiency between fomites and skin. Importantly, we only observed a transfer between fomites and skin using a large initial virus titer sample (10^6^ infectious virus particles) on the tested surfaces, while lower initial virus titer stocks (10^4^ infectious virus particles) were not effectively transferred.

Overall, our results point to a low risk of SARS-CoV-2 transmission by coins and banknotes and the tendency to prefer contactless payment over cash during the pandemic seems unnecessary.

## Materials and methods

### Preparation of test virus suspension

For preparation of SARS-CoV-2 test virus suspension, Vero E6 cells were seeded in a 75 cm^2^ flasks at 2×10^6^ cells in Dulbecco’s Modified Eagle’s Medium (DMEM, supplemented with 10 % (v/v) fetal calf serum (FCS), 1 % non-essential amino acids, 100 IU/mL penicillin, 100 µg/mL streptomycin and 2 mM L-Glutamine). The monolayer was inoculated with the hCoV-19/Germany/BY-Bochum-1/2020 (GISAID accession ID: EPI_ISL_1118929). After 3 days and upon visible cytopathic effect the supernatant was harvested by centrifugation at 1500 rpm for 5 min at room temperature, aliquoted and stored at −80 °C until further usage.

For preparation of BCoV virus suspension, U373 cells were cultivated in a 75 cm^2^ flask with in Minimum Essential Medium Eagle (EMEM) supplemented with L-glutamine, non-essential amino acids and sodium pyruvate and 10 % FCS. Before virus infection, cells were washed two times with phosphate buffered saline (PBS), incubated for 3 h with serum-free EMEM and were washed once with EMEM supplemented with trypsin. For virus production, BCoV strain L9 (NCBI: txid11130) was added to the prepared monolayer. After an incubation period of 24 to 48 hours cells were lysed by a rapid freeze/thaw cycle followed by a low speed centrifugation in order to sediment cell debris. After aliquoting of the supernatant, test virus suspension was stored at −80 °C. Nine volumes of test virus suspension were mixed with one volume of interfering substance solution [0.3 g/L bovine serum albumin (BSA) in PBS according to EN 16777, section 5.2.2.8]. The tests were performed with two different virus concentrations, i.e. a titer of approximately 10^4^ 50% tissue culture infectious dose per milliliter (TCID_50_/mL) and a titer of 10^6^ TCID_50_/mL.

### Preparation of specimens

Prior to use regular 5-, 10-cent and 1-euro coins were dipped in a bath containing 70 % (v/v) ethanol for 5 min. The 10-and 50-euro banknotes (provided by the European Central Bank) and PVC plates [with PUR (polyurethane) surface coating 20 x 50 cm (VAH e.V.), precleaned with 70.0 % propan-1-ol or ethanol] were cut into pieces of 2 x 2 cm. Banknotes were UV irradiated before the tests. Stainless steel discs (2 cm diameter discs) with Grade 2 B finish on both sides (article no. 4174-3000, GK Formblech GmbH, Berlin, Germany) served as reference control. Prior to use the discs were decontaminated with 5 % (v/v) Decon 90 for 60 minutes and 70 % (v/v) propan-2-ol for 15 min. Subsequently, the discs were rinsed with distilled water sterilized by autoclaving (steam sterilization).

### Inactivation assays and controls

For stability testing, specimens were placed aseptically in a Petri dish and inoculated with 50 μL of the virus inoculum [5 × 10 µL drops, i.e. four in every corner and one in the middle of the square (Fig. 3)]. After visible drying of the inoculum, the petri dishes were closed and the specimens were incubated until the end of the appropriate exposure time (up to 7 days). After the respective time, the specimens were transferred to 2 mL cell culture medium (without FCS) in a 25 mL container and vortexed for 60 seconds to resuspend the virus. Directly after elution, series of ten-fold dilutions of the eluate in ice-cold maintenance medium were prepared and inoculated on cell culture.

Fifteen and 30 minutes, 1, 2, 7, and 24 hours and 2, 3, 5 and 7 days were chosen as application times. Eluates were retained after appropriate drying times and residual infectivity was determined.

The initial virus titer (VIC) was determined by addition of 50 µL of the virus inoculum directly to 2 mL cell culture medium without any desiccation.

### Touch transfer test

For the touch transfer test with BCoV, three test persons simulated the transfer by pressing a finger shortly on the dried inoculum on the respective carriers followed by rubbing once with pressure over the carrier. Three other test persons simulated the transfer by a fingerprint of 5 seconds on the dried inoculum on the different carriers. Each test person performed the transfer test separately with the two different virus concentrations (10^4^ TCID50/mL and 10^6^ TCID50/mL) with 8 fingers each. For each test person and virus concentration, two fingers were used for virus transfer without drying of the inoculum. The transfer procedure was the same as with the dried inoculum.

The amount of transferred virus to the fingers was obtained by dipping and rubbing each finger in turn for one minute on the base of a Petri dish containing 2 mL cell culture medium without FCS as sample fluid. For each finger a separate dish was used. The eluates were transferred in a 25 mL container. Directly after elution, series of ten-fold dilutions of the eluate in ice-cold maintenance medium were prepared and inoculated on cell culture. The initial virus titer (VIC) was determined by addition of 50 μL of the virus inoculum directly to 2 mL cell culture medium without any drying. Furthermore, a cell control (only addition of medium) was incorporated.

For the touch transfer test of SARS-CoV-2, one person performed all assays due to BSL3 restrictions. To mimic the texture and nature of human fingertips, we used VITRO-SKIN (IMS Florida Skincare Testing, FL, USA), an artificial skin substitute, placed in a plastic frame was used (Fig. 3). After printing or rubbing as described above, the complete artificial skin was released from the frame and transferred into a 25 mL container with serum-free cell culture medium and vortexed for 60 s.

### Determination of infectivity

Infectivity was determined by means of end point dilution titration using the microtiter process. For this, samples were immediately diluted at the end of the exposure time with ice-cold EMEM containing trypsin and 100 μL of each dilution were placed in 6 or 8 wells of a sterile polystyrene flat-bottomed plate with a preformed U373 (BCoV) or Vero 6 (SARS-CoV-2) monolayer. Before addition of virus, cells were washed twice with EMEM (U373) or DMEM (Vero 6) and incubated for 3 h with 100 µL EMEM (U373) or DMEM (Vero 6) with trypsin. After 3 d or 6 d incubation at 37 °C in a CO_2_-atmosphere (5.0 % CO_2_-content), cultures were observed for cytopathic effects. TCID_50_/mL was calculated according to the method of Spearman and Kärber [16].

### Fitting of virus titer decay

To model the decay in virus titer, we implemented a Weibull distribution fit in GraphPad Prism version 9.0.2 for Windows (GraphPad Software, San Diego, California USA, www.graphpad.com)

### Calculation of the reduction factor

The loss in virus titer by desiccation was calculated by subtracting the titer on the different carriers after desiccation from the titer of the initial virus control. The amount of transferred virus (TCID_50_/mL) from the different carriers to the fingers was also calculated with the method of Spearman and Kärber [16].

## Results

### Stability of BCoV on euro banknotes

To examine the stability of coronaviruses on banknotes we first used bovine coronavirus (BCoV), which can be cultivated under lower biosafety levels and has been used as a surrogate virus for inactivation studies replacing the highly pathogenic MERS-CoV and SARS-CoV [17]. All euro banknotes are made of pure cotton fiber. To protect the surface of banknotes with smaller denomination and prolong circulation life, 5 € and 10 € banknotes are coated with a varnish applied after printing [18]. To account for the effect of this varnish on surface stability of BCoV over time, we assessed residual infectivity from pieces of 10 € and 50 € banknotes for 7 h, 24 h and subsequently every 24 – 48 h up to 7 days (Fig. 1A). The initial virus concentration of 4.3 × 10^6^ TCID_50_/mL declined to 1.84 × 10^4^ TCID_50_/mL on 10 € banknotes and 9.25 × 10^4^ TCID_50_/mL on 50 € banknotes after 7 h desiccation. To quantitatively compare this early loss of titer on the different surfaces, we employed a fitted Weibull distribution model to estimate initial decay rates and the modelled time to lower limit of quantification (Fig. 1B, Table 1). For both banknotes we observed shorter initial decay (2.75 h on 50€ and 6.45 h on 10€) as compared to the steel disc (49.62 h) (Fig. 1B, Table 1). Following the strong initial decay, we were able to detect low amounts of infectious virus after 120 h (50€) and 168 h (10€) respectively (Fig. 1A), which is very much in line with the observed times in the model of 175.62 h for 50€. and 216.31 h for 10€ notes (Fig. 1B and Table 1). In contrast, on steel discs a more continuous decay was observed and infectious virus could be recovered up to 120 h (Fig. 1A), and 229.73 h for the fitted model, respectively (Fig. 1B, Table 1).

**Table 1:** Initial decay time and time to reach LLOQ calculated from modelled curves.

|  |  | SARS-CoV-2 |  | BCoV |  |
| --- | --- | --- | --- | --- | --- |
|  | material | initial decay [h] | time to LLOQ [h] | initial decay [h] | time to LLOQ [h] |
| notes | 50 euro |  |  | 2.8 | 175.6 |
|  | 10 euro | 6.1 | 85.7 | 6.5 | 216.3 |
| coins | 1 euro | 2.2 | 28.4 |  |  |
|  | 10 cent | 0.8 | 2.3 |  |  |
|  | 5 cent | 0.2 | 0.6 |  |  |
| control | steel disc | 20.6 | 158.8 | 53.5 | 240.2 |

**Figure 1:**
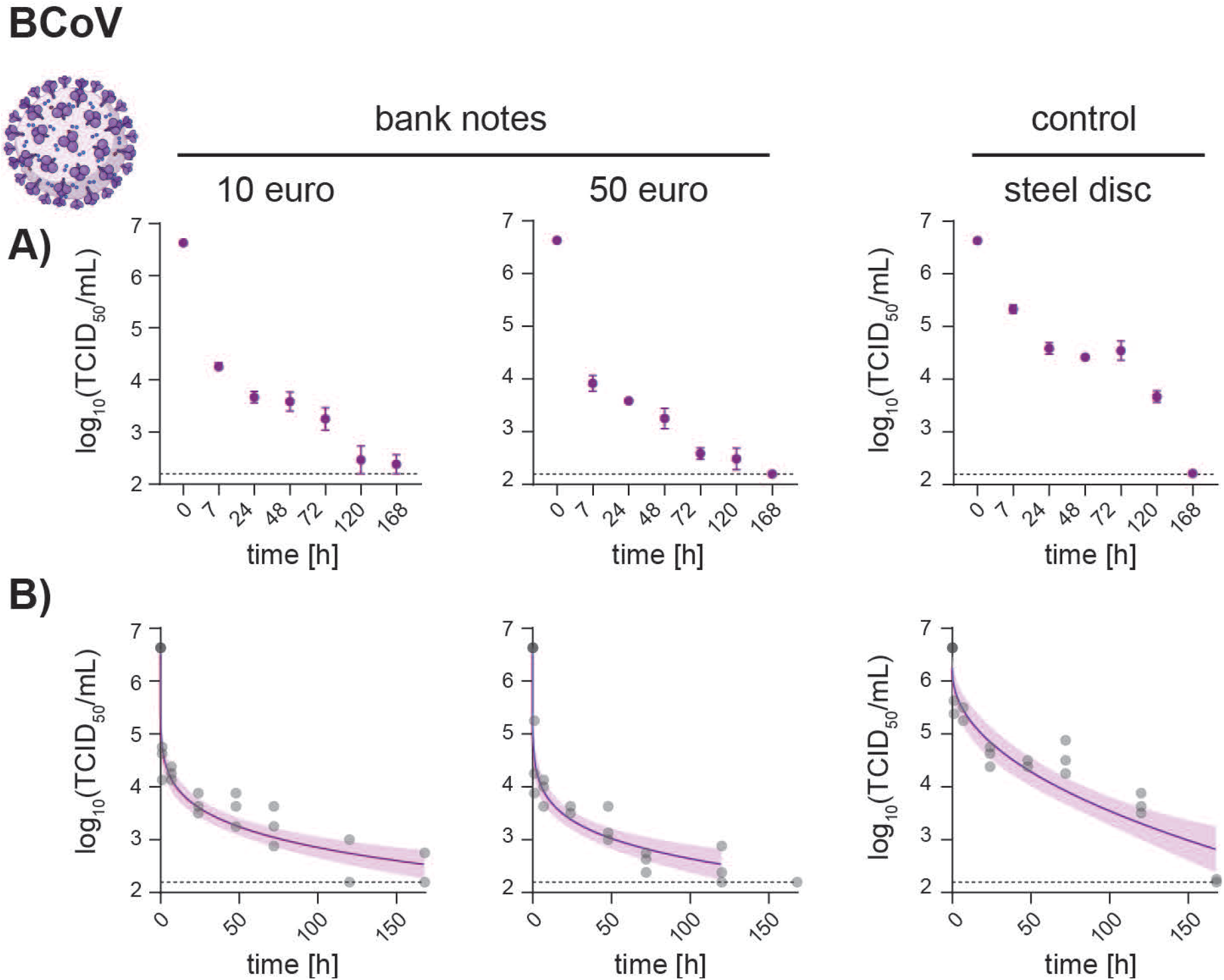
Stability of BCoV on banknotes and steel discs. BCoV stock solution was applied on 2 cm × 2 cm pieces of 10 € or 50 € banknotes and recovered after the indicated times. Residual titer was assessed via limiting dilution assay. **A)** Infectious BCoV recovered, displayed as raw TCID_50_/mL (y-axes) over time (categorical x-axes). Dots indicate mean values of three independent experiments with standard deviation, lower limit of quantification is shown as dashed line. **B)** Recovered BCoV displayed as TCID_50_/mL (y-axes) over time (continuous x-axes). Dots represent individual biological experiments, purple lines and areas display the course of the Weibull distribution fitted data and 95% confidence interval, lower limit of quantification is shown as dashed line. Virus particles created with BioRender.com.

### Stability of SARS-CoV-2 on euro banknotes and coins

We next examined the surface stability of infectious SARS-CoV-2 on 10 € banknotes, different coins (1 €, 10 cents, 5 cent) and stainless-steel discs for up to 7 days using an initial virus concentration of 1.36 – 2.0 × 10^6^ TCID_50_/mL (Fig. 2). On 10 € banknotes and 1 € coins, the initial virus concentration declined to 2.32 × 10^4^ TCID_50_/mL and 1.79 × 10^4^ TCID_50_/mL, respectively, after 1.25 h, corresponding to an estimated initial decay time of 6.07 h and 2.21 h (Fig. 2B, Table 1). No infectious virus could be recovered after 72 h and 48 h (Fig. 2A) matching 85.67 h and 28.43 h survival time (Fig. 2B, Table 1). In contrast, on 10 cent and 5 cent coins the initial virus concentration declined to 5.96 × 10^4^ TCID_50_/mL and 3.86 × 10^1^ TCID_50_/mL, respectively, within 30 min. Initial decay rates were calculated as 49.8 min (10 cent) and 12 min (5 cent) (Fig. 2B, Table 1). Importantly, from 10 cent coins no infectious virus could be recovered after 6 h, while for 5 cent coins infectivity was completely lost after 1 h (Fig. 2A), as reflected by 2.28 h and 33 min survival time for SARS-CoV-2 on 10 cent and 5 cent coins (Fig. 2B, Table 1). In contrast, on stainless-steel discs, which served as reference material, initial decay and time to reach background levels were comparable to BCoV with 20.59 h and 158.83 h, respectively (Fig. 2B, Table 1). Virus titers declined more evenly until no infectious virus could be recovered after 120 h (Fig. 2A).

**Figure 2:**
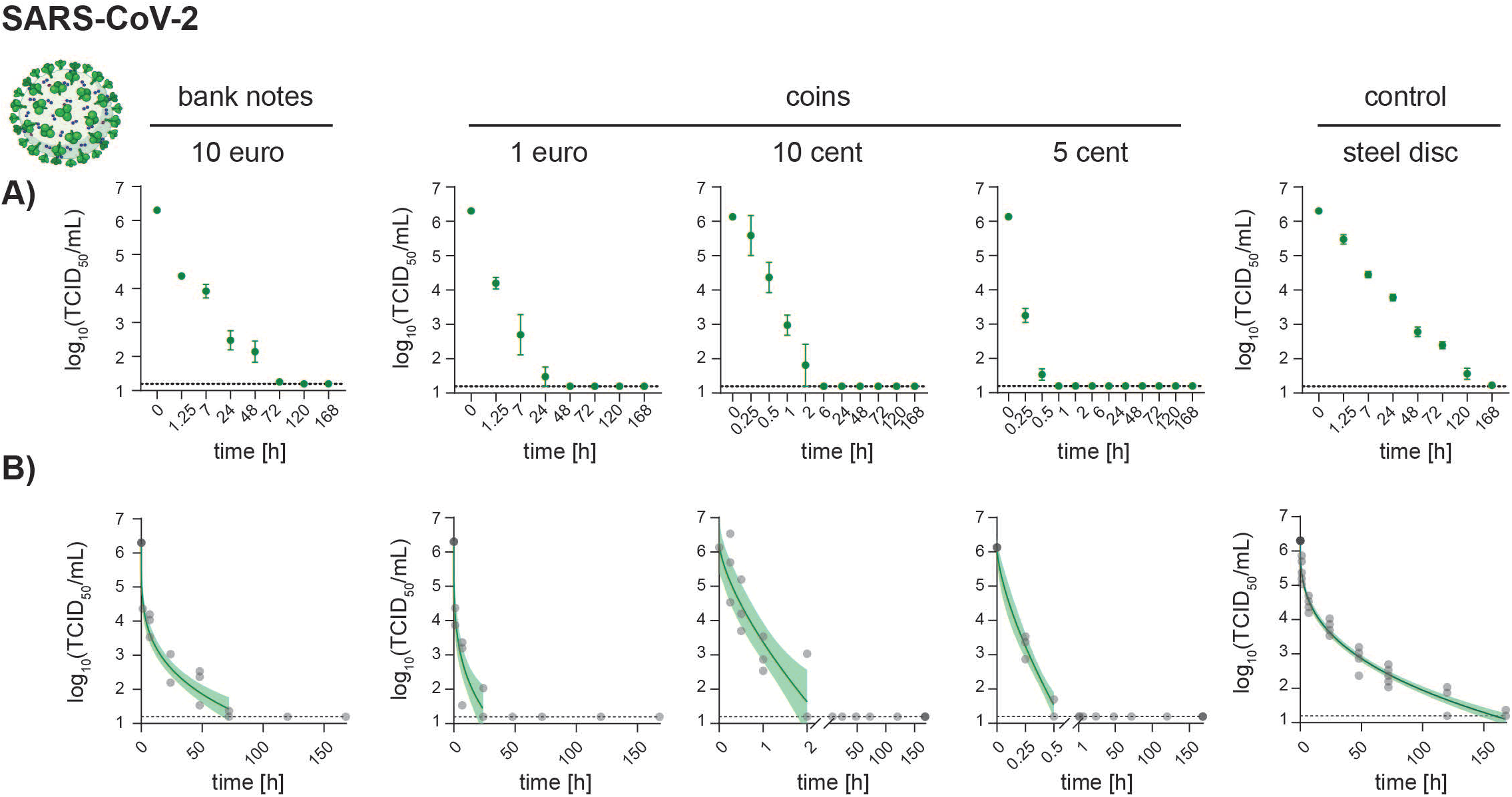
Stability of SARS-CoV-2 on banknotes, coins and steel discs. SARS-CoV-2 stock solution was applied on 2 cm × 2 cm pieces of 10 € banknotes, 1 €, 10 cent and 1 cent coins and recovered after the indicated times. Residual titer was assessed via limiting dilution assay. Humidity and temperature during experiments was logged (32% - 43% RH, 22.4 °C – 23.2 °C) **A)** Infectious SARS-CoV-2 recovered, displayed as raw TCID_50_/mL (y-axes) over time (categorical x-axes). Dots indicate mean values of three independent experiments with standard deviation, lower limit of quantification is shown as dashed line. **B)** Recovered SARS-CoV-2 displayed as TCID_50_/mL (y-axes) over time (continuous x-axes). Dots represent individual biological experiments, green lines and areas display the course of the Weibull distribution fitted data and 95% confidence interval, lower limit of quantification is shown as dashed line. Virus particles created with BioRender.com.

### Development of a touch transfer assay to study virus transfer between cash and finger pads

Experiments performed under controlled laboratory conditions demonstrated the persistence of SARS-CoV-2 on inanimate surfaces for days and consequently implied the risk of viral transmission via contaminated objects [5,19]. However, to develop more refined models to assess the risk of fomites-based transmission of SARS-CoV-2, quantitative measurements of the transfer efficiency of infectious virus between skin and surfaces are required. To address these limitations, we developed a touch transfer assay to study the transfer of infectious BCoV and SARS-CoV-2 between finger pads and different fomites (Fig. 3). Briefly, virus suspensions were placed on different surfaces (pieces of 10 € banknotes, 10 cent coins, pieces of PVC to mimic the surface of credit cards and stainless-steel discs as reference material). Afterwards, the wet inoculum or the dried suspension was touched by “printing” or “rubbing” using finger pads (BCoV) or an artificial skin fabric (SARS-CoV-2) (Fig. 3). Subsequently, infectious viruses were recovered by dipping and rubbing each fingertip in turn for one minute on the base of a Petri dish containing 2 mL of EMEM cell culture medium (BCoV) or, in case of the artificial skin, by directly placing it into a container with cold DMEM (SARS-CoV-2). The resulting suspension was serially diluted to determine TCID_50_/mL values of the remaining infectious virus.

**Figure 3:**
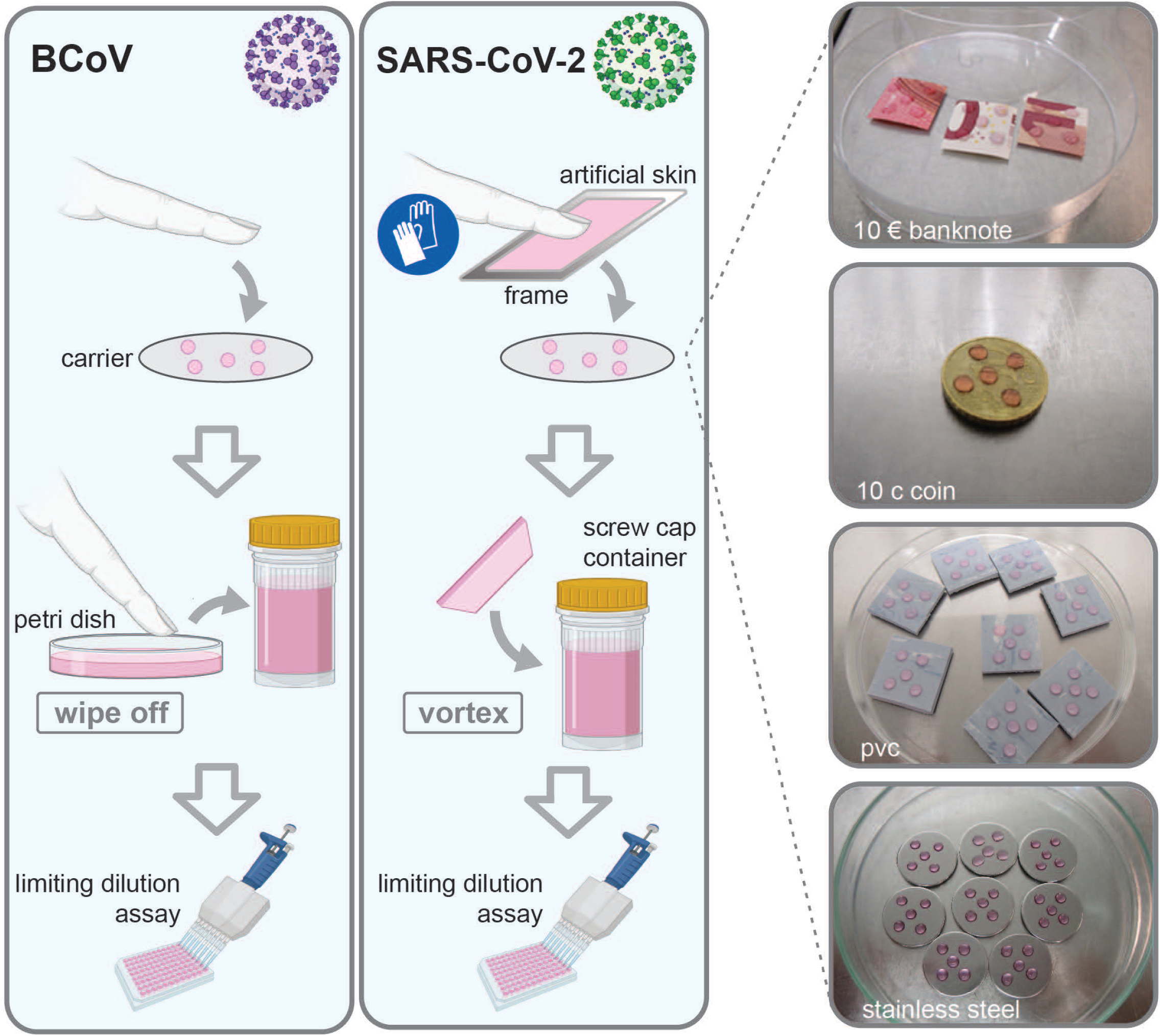
Touch transfer assay setup. To study the transfer of infectious BCoV and SARS-CoV2 between finger pads and different fomites, 50 µL virus suspensions are placed on different surfaces (pieces of 10 € banknotes, 10 cent coins, pieces of PVC to mimic the surface of credit cards and stainless-steel discs as reference) in 10 µL spots. Afterwards, the wet inoculum or the dried suspension is touched by “printing” or “rubbing” using finger pads (BCoV) or an artificial skin fabric (SARS-CoV-2). Subsequently, infectious virus was recovered by rubbing the fingertip on the bottom of a petri dish filled with respective culture media or in case of the artificial skin directly transferred into a container. The resulting suspension is serially diluted to determine TCID_50_/mL values of the remaining infectious virus. Virus particles created with BioRender.com.

### Transferability of BCoV from banknotes, coins and PVC to fingertips

Using this newly developed touch transfer assay, we examined the transmission of BCoV from different surfaces, i.e. pieces of 10 € banknotes, 10 cent coins, pieces of PVC and stainless-steel discs as reference material, to fingertips. Surfaces were inoculated with either a high (∼ 1 × 10^6^ TCID_50_/mL) or low (∼ 1 × 10^4^ TCID_50_/mL) viral titer to represent different degrees of surface contamination. Virus transfer was assessed directly following application to fomites (wet) or after ∼ 1 h until completely dried (dry) by either pressing (print) or rubbing (rub) the fingertip onto the surface. Initial virus (input) was determined by applying the fomites directly to the medium container. For a high initial titer and direct surface contact, we observed a maximum of a 0.6 log_10_ reduction for the 10-cent coin, while lower reduction factors were observed for the other surfaces (Fig. 4A). In case of drying the initial inoculum followed by a fingerprint, we observed a 2.1 log_10_ reduction on a 10 € banknote, while lower reduction factors were observed for the other surfaces. For a low initial titer and direct surface contact, we observed the highest reduction on the stainless-steel carrier (1.2 log_10_ reduction). In case of drying the initial inoculum followed by a fingerprint, we observed a 0.8 log_10_ reduction on a 10-cent coin. Importantly, no infectious virus could be recovered from the 10 € banknote under these conditions.

**Figure 4:**
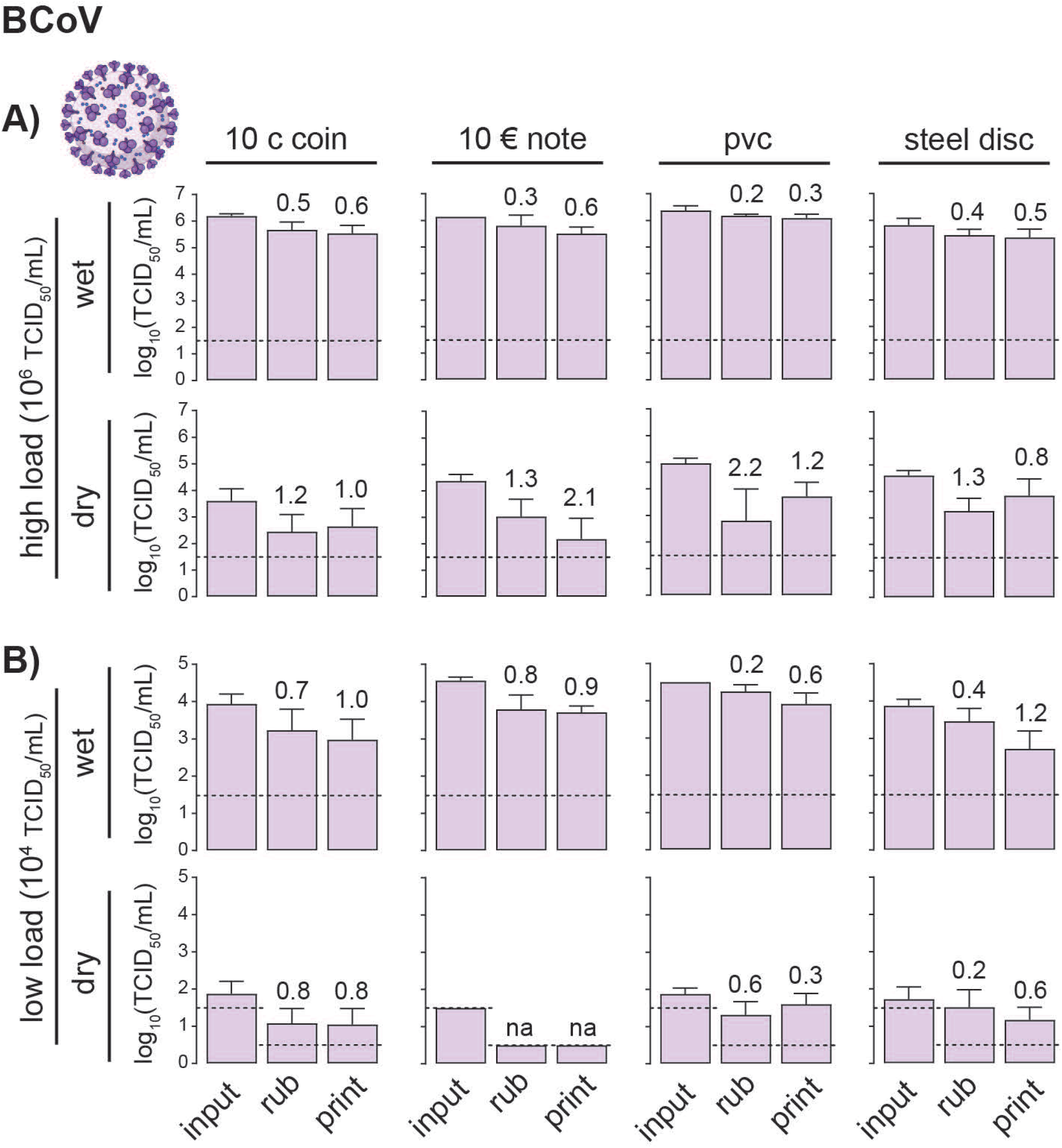
Transferability of BCoV from cash fomites to fingertips. Bars depict titer of input virus suspension and recovered infectious virus from different cash fomites, i.e. 10 cent coin, 10 € banknote, pvc and steel disc carrier in four different scenarios. **A)** High initial input titer (∼10^6^ TCID_50_/mL) wet, when directly touch after application and dry, when transferred after visual desiccation and **B)** low initial input titer (∼10^4^ TCID_50_/mL), wet and dry. Each scenario was performed by three test persons using eight fingers each. Numbers above bars indicate reduction factor, lower limit of quantification is shown as dashed line.

### Transferability of SARS-CoV-2 from banknotes, coins and pvc to skin

Next, we examined the transmission of infectious SARS-CoV-2 from surfaces to fingertips. Surfaces were inoculated with either a high (∼ 1 × 10^6^ TCID_50_/mL) or low (∼ 1 ×10^4^ TCID_50_/mL) titer to represent different degrees of surface contamination. As described before, virus transfer was assessed directly following inoculation (wet) or after drying either by printing (print) or rubbing (rub). For a high initial titer and direct surface contact, we observed a maximum of a 1 log_10_ reduction for the 10-cent coin, while lower reduction factors were observed for the other surfaces (Fig. 5A). Drying of the initial inoculum led to ∼ 1 log loss in virus titer. In the dried state, less virus was transferred and could be recovered, e.g. by fingerprint we observed a 3.0 log_10_ reduction on the 10-cent coin, while lower reduction factors were observed for the other surfaces. For a low initial titer and direct surface contact, we observed the highest reduction on the 10 € banknote (0.7 log_10_ reduction). In case of drying the initial inoculum followed by a fingerprint we observed a reduction of the initial inoculum close/under the limit of detection and only from the PVC very low (2.19× 10^1^ TCID_50_/mL) amounts of infectious virus could be recovered (Fig. 5B).

**Figure 5:**
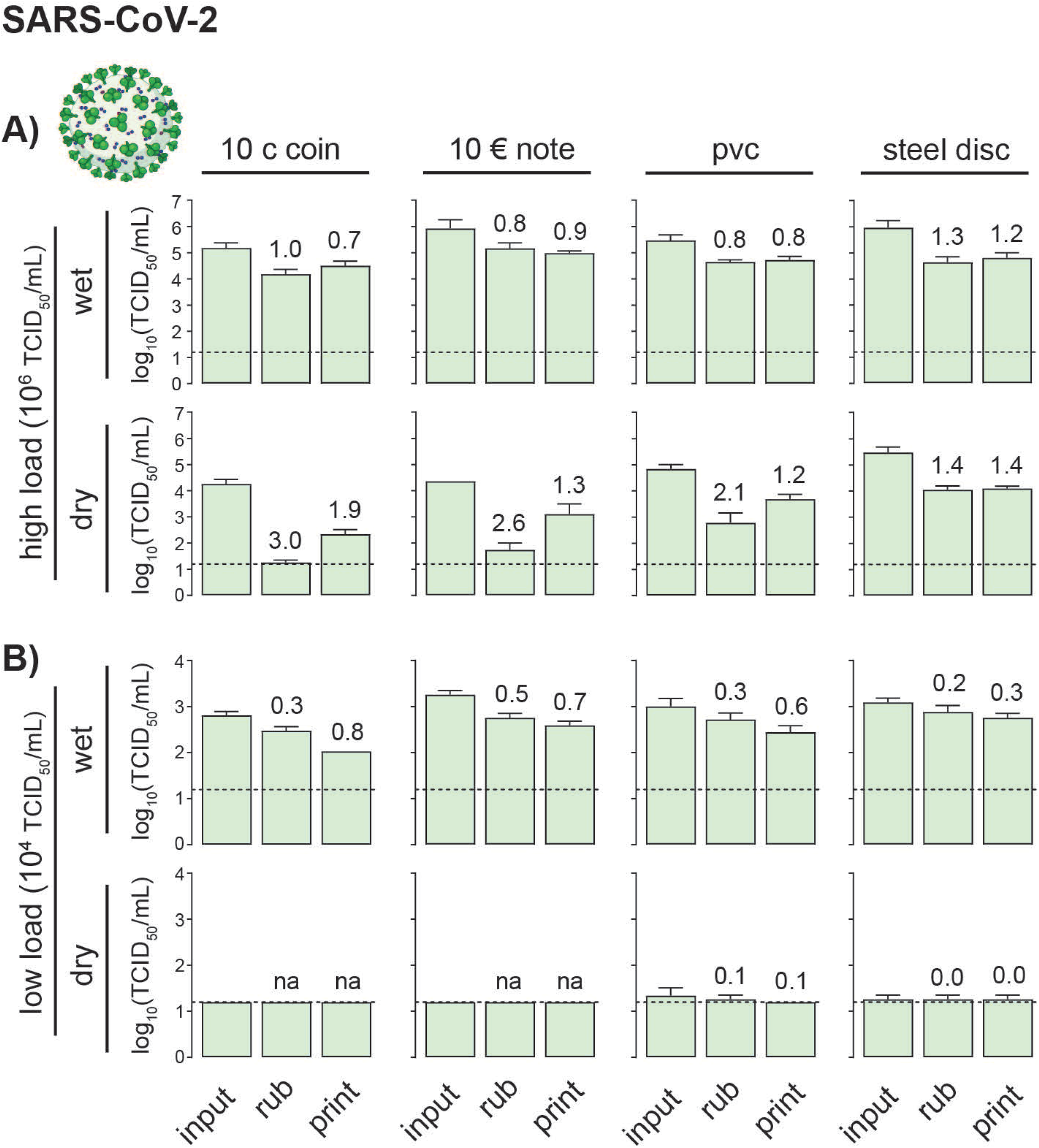
Transferability of SARS-CoV-2 from cash fomites to fingertips. Bars depict titer of input virus suspension and recovered infectious virus from different cash fomites, i.e. 10 cent coin, 10 € banknote, pvc and steel disc carrier in four different scenarios; mean ± SD. Humidity and temperature during experiments was logged (32% - 43% RH, 22.4 °C – 23.2 °C) **A)** High initial input titer (∼10^6^ TCID_50_/mL) wet, when directly touch after application and dry, when transferred after visual desiccation and **B)** low initial input titer (∼10^4^ TCID_50_/mL), wet and dry. Numbers above bars indicate reduction factor, lower limit of quantification is shown as dashed line. Virus particles created with BioRender.com.

## 4. Discussion

Human-to-human transmission of SARS-CoV-2 occurs primarily by respiratory aerosols or droplets and subsequent contact to nasal, oral, or ocular mucosal membranes. Evidence-to-date further suggests that fomite transmission is possible for SARS-CoV-2 [5,19], however, the importance of this route in healthcare and public settings remains controversial [12,13,20]. Fomite-based transmission contributes to the spread of other common respiratory pathogens [21,22]. Consequently, paper currency and coins have been suspected as a potential transmission vehicle for various pathogens, including SARS-CoV-2 [10,11,23]. Although infectious viruses have not been directly detected on banknotes or coins, the potential for their transmission has been highlighted by the observation that human influenza viruses were able to persist and remain infectious for several days when they were deposited on banknotes [24]. Furthermore, many other viruses, (i.e. Adenoviruses, Rotaviruses) are stable in the environment and exhibit high infectivity and, thus, could possibly be transferred by banknotes and coins [25]. In agreement with previous reports we found that high titers of SARS-CoV-2 and its surrogate BCoV, after an initial loss of infectivity, remained infectious for days under laboratory conditions on banknotes and coins (Table 1, Fig. 1 and 2) [19,26]. The initial loss of infectivity was higher on coins and banknotes, irrespective of protective varnish, when compared to stainless steel, indicating faster desiccation due to liquid absorption (banknotes) or antiviral surface properties (e.g. copper in coins). Both BCoV and SARS-CoV-2 displayed highly comparable levels of virus transfer and stability among the different conditions (Fig. 6), implying that BCoV is also a suitable surrogate virus to model surface transmission of SARS-CoV-2.

**Figure 6:**
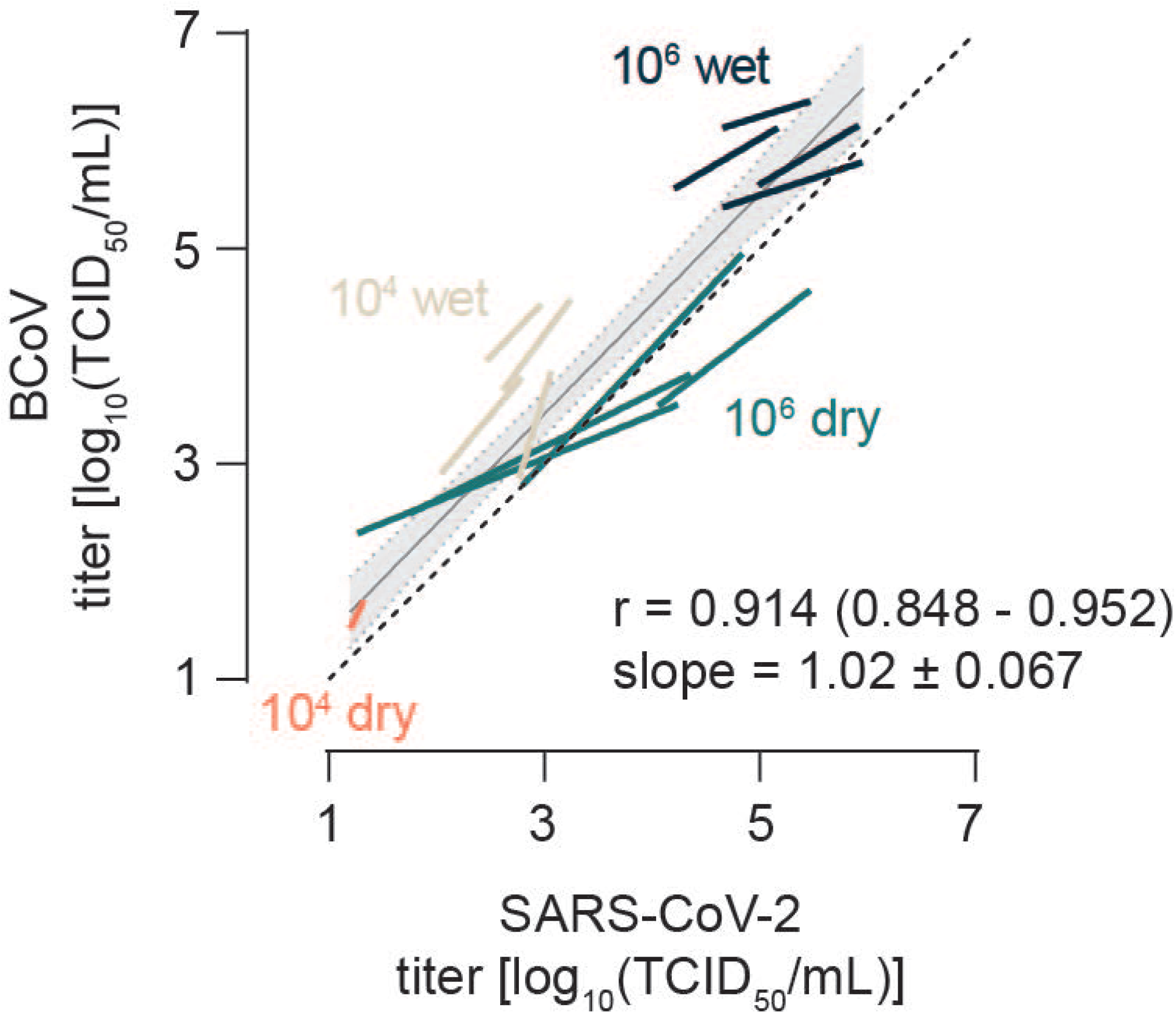
Suitability of BCoV as surrogate for SARS-CoV-2 in touch transfer studies. Titers of recovered infectious virus were matched between BCoV and SARS-CoV-2 for each scenario and linear regression curves calculated for input, rub and print. Gray line and area represent the overall linear regression and 95% confidence interval of all matched data points, dashed line depicts perfect correlation.

Decay of SARS-CoV2 is likely determined by a combination of the initial amount of infectious virus deposited on a given surface and other environmental parameters (temperature, humidity, light and UV conditions). Furthermore, persistence of pathogens in the environment represents only the first requirement for self-inoculation via contaminated fingers. However, the possibility of fingerprint transmission has quantitatively been examined only in the context of bacteria [27]. Using a newly developed virus touch transfer assay, we observed transfer of BCoV and SARS-CoV-2 between fomites and skin using a high initial virus titer (∼10^6^ infectious virus particles). This transfer was more efficient for the wet inoculum, while visual desiccation on the one hand resulted in reduction of the titer as outlined above, as well as less efficient mobilization of the viral particles, reflected by higher reduction factors. Consequently, lower viral burdens (∼10^4^ infectious virus particles) mimicking real life contamination events more realistic, as observed for influenza viruses in aerosol particles from human coughs [13,28], were not effectively transferred (Fig. 4 and 5). Recent studies estimated a minimal infectious dose of SARS-CoV-2 in the range of 3 × 10^2^ to 2 × 10^3^ viral particles [29]. Overall, our results point to a low risk of SARS-CoV-2 transmission by coins and banknotes and the rush to abandon cash during the pandemic seems unnecessary.

Given that cash is typically stored securely in wallets and purses, the risk of direct contamination through exhaled droplets and aerosols seems much lower than constantly exposed surfaces (e.g. doorbell, shopping carts). The role of a contagious person contaminating banknotes and coins afresh when handing over, needs to be addressed in future studies. Current government regulations to wear masks minimize the spread of exhaled droplets and aerosols, and in combination with good hand hygiene also mitigate the risk of transmission via contaminated surfaces. Still, contamination of cash is most likely to occur indirectly by transfer from the hands of an infected person or finger contact with a contaminated surface. However, any contamination by these routes would likely result in a much lower degree of surface contamination than by direct contamination as investigated in this study. Consequently, the overall chance of transmission of SARS-CoV-2 through banknotes, coins and credit/debit cards seems low since a timely order of specific events – sufficient viable virus deposited on a surface, survival of the virus until the surface is touched, and transfer of an infectious dose of virus – is required.

## Acknowledgments

We thank Victor M Corman and Terry C Jones from Charité Universitätsmedizin Berlin, Germany for sequencing of the SARS-CoV-2 strain used in this study. Moreover, we would like to thank all members of the Department of Molecular & Medical Virology at the Ruhr University Bochum for support and fruitful discussion.

